# Production of common carp donor-derived offspring from goldfish surrogate broodstock

**DOI:** 10.1101/2020.08.11.245803

**Authors:** Roman Franěk, Vojtěch Kašpar, David Gela, Martin Pšenička

## Abstract

**Background:** Common carp is the fourth most-produced species in worldwide aquaculture. Significant efforts are invested in breeding and preservation of genetic integrity of this important species. However, maintaining carp gene bank *in situ* can be considered as demanding due to its big body size. Recent progress in reproductive biotechnologies in fish allows improving some unfavourable characteristics of a target species using surrogate reproduction. Germ stem cells (gamete precursors) from one species are transplanted into different surrogate species with small body size. After maturation, surrogates are producing donor-derived progeny. Efficient protocols for cryopreservation of carp male and female germ stem cells have been developed lately. Thus, the next logical goal was to assess the potential of goldfish surrogate to produce donor-derived gametes of common carp after intraperitoneal transplantation of testicular cells.

**Results:** High transplantation success was achieved when 44% of the surviving goldfish produced pure donor-derived gametes of common carp. More importantly, both viable eggs and sperm giving rise to pure common carp progeny were produced, witnessing sustainability of the presented method. Donor-derived identity of the offspring was confirmed by genotyping and typical phenotype corresponding to the donor species. Reproductive performance of chimeras was similar to goldfish controls. Assessment of gamete characteristics showed that the size of donor-derived eggs is between control carp and goldfish eggs. Interestingly, flagellum length in donor-derived spermatozoa was comparable to common carp flagellum and significantly shorter than goldfish flagellum.

**Conclusions:** In this study, we succeeded in the production of pure common carp progeny from surrogate goldfish recipients transplanted intraperitoneally by testicular germ cells. Here we reported production of viable eggs between most distant species up to date. Good reproductive performance of goldfish germline chimeras gives a promising prospect for further analysis about the long-term reproductive performance of surrogates, recovery of cryopreserved germ cells or production of monosex stocks. Presented technology is ready to ease needs for carp breeds preservation and their recovery using many times smaller goldfish surrogates.

## Background

Common carp (*Cyprinus carpio*) is the fourth most-produced fish species in aquaculture worldwide, mainly produced in Europe and Asia [1]. Therefore, a significant amount of work has been made to establish breeding programs and improve common carp performance in aquaculture [2–4]. Breeding programs are accompanied with efforts to develop methods for preservation of current purebreds because their F1 hybrids are used aquaculture. Appropriate measures are employed to protect the unique genetic resources of parental breeds. Common carp broodstock is maintained in live gene banks mostly [5,6]. However, breeds might be under risk of genetic contamination, loss of genetic variability or unexpected environmental/technical accidents [6]. Therefore, cryopreservation management is employed to secure at least paternal part of live gene banks [7,8]. Unfortunately, mature fish oocytes cannot be cryopreserved efficiently, thus maternal genetic resources are more threatened and alternative managing strategies are necessary. So far, the only realistic alternative on how to tackle this issue in fish is the utilization of germ stem cells (GSCs) manipulation techniques for surrogate reproduction. Germ stem cells are a population of cells found in the gonads having the capability to differentiate into the gametes, but also maintain stemness via self-renewal. In that case, GSCs can be obtained through the whole lifetime of the fish as embryonic primordial germ cells and later on as differentiated spermatogonia or oogonia. These GSCs can be isolated and transplanted into the body of surrogate host by various methods. Host’s genital ridge is in turn colonized by transplanted GSCs with subsequent proliferation and production of donor-derived gametes [9–11]. Of the utmost importance, GSCs have a high level of sexual plasticity. Transplanted male GSCs can transdifferentiate into female GSCs giving rise to viable eggs and vice versa in the environment the host’s gonadal environment. Sexual plasticity of GSCs has immense potential when both sperm and eggs can be produced even from a single donor. Transplantation can also facilitate to more advanced application. Monosex stocks can be produced from donors with homogametic sex chromosomes [12] without the need to use challenging techniques of uniparental inheritance induction or hormonal treatment having environmental concerns [13]. Interspecific surrogacy is also believed to be a convenient tool for the management of endangered [14–16] or important aquaculture species [17–20]. In both cases, interspecific transplantation can ameliorate unfavourable characteristics of target species such as long maturation, big body size and consequently high space and costs requirements associated with their maintenance [21] or in overall problematic reproduction in captivity [22,23]. However, production of donor-derived seed from iconic species such as long maturing sturgeons [16,21,24,25] or bluefin tuna [22,23,26] has not been achieved yet, despite significant and long-lasting efforts.

Up to date, most of the surrogacy studies assessed the suitability of chosen host and transplantation procedure, because both are vital prerequisites conditioning successful gamete production. Numerous studies described successful donor-derived gamete production after intraspecific transplantation. Viable egg production from interspecific surrogates has been reported only in three species so far, suggesting certain difficulties in terms of evolutionary distance between two different species which may cause some cell incompatibilities. On the other hand, donor-derived sperm production, especially in connection with intrapapillary transplantation, seems to have no boundaries. This can be outlined by successful surrogate sperm production in species differing in taxonomic class [27] with estimated divergence time 230 million years ago (mya). However, currently most distant successful production of viable donor-derived eggs and sperm have been achieved by pioneering study on surrogacy between masu salmon recipients and rainbow trout donor [28] having divergence time 14.2 mya. Therefore, we emphasize that egg production from interspecific surrogates with some superior characteristics should be the ultimate aim in GSCs manipulation technologies, in order to fully utilise potential of this reproductive biotechnology.

Protocols for cryopreservation of common carp male [29] and female [30] gonadal tissue containing a population of pluripotent GSCs have been developed recently in our laboratory as the first step towards novel reproductive biotechnologies development in carp. Subsequently, we confirmed that goldfish can accept intraperitoneally transplanted GSCs of common carp, thus giving a positive outlook to establish sustainable surrogacy between carp and goldfish. In this study, we focused to achieve production of common carp gametes via smaller goldfish surrogates and to evaluate their reproductive performance, characterise produced gametes and to confirm whether goldfish germline chimeras are capable to produce surrogate gametes repeatedly. Presented technology serves as a vital alternative for *in vivo* preservation of genetic resources in one of the most important fish species worldwide. To the best of our knowledge, there is no report on successful surrogacy on the species with similar aquaculture significance such common carp has.

## Methods

Experimental protocol was approved by Ministry of Agriculture of the Czech Republic (reference number: 55187/2016-MZE-17214). The methodological protocol of the current study was approved by the expert committee of the Institutional Animal Care and Use Committee of the University of South Bohemia in České Budějovice, Faculty of Fisheries and Protection of Waters in Vodňany according to the law on the protection of animals against cruelty (Act no. 246/1992 Coll., ref. number 16OZ19179/2016-17214). The study did not involve endangered or protected species. Vojtěch Kašpar (CZ01652), David Gela (CZ01672) and Martin Pšenička (CZ 00673) are qualified to manage and conduct experiments involving animals according to section 15d paragraph 3 of Act no. 246/1992 Coll.

### Chemicals

Unless stated otherwise, all chemicals were purchased from Sigma Aldrich (St. Louis, MO, USA), catalogue numbers/names of the used chemicals are presented in the brackets at the first mention. Brine shrimp eggs were obtained from Ocean Nutrition Europe (Belgium), dry diets for progeny ongrowing were Scarlet and Pre Growe from Alltech Coppens (The Netherlands).

### Recipient production and transplantation

Goldfish (*Carassius auratus)* broodstock for recipients production originated from the same resource as in our previous studies [29,30]. Goldfish gametes were obtained after hormonal stimulation. Aceton dried carp pituitary (Rybářství Klatovy s.r.o.) was minced and dissolved at 0.9% physiological. Suspension was injected intraperitoneally at two doses for females, first dose of 0.5 mg per kg of body weight (mg/kg), second dose of 2.5 mg/kg (12h after the first dose). Males were injected with single dose of 1.5 mg/kg. Gametes were obtained approximately 24h after the first dose administration. Fertilization was done with pooled eggs from females (n 5) and pooled sperm from males (n 10). Water-activated eggs were gently stirred on glass Petri dishes and allowed to stick on the surface. Embryos were not dechorionated.

Developing embryos at one to four-cell stage were injected with 100 mM solution of antisense *dead end* morpholino Gene Tools LLC (Philomath, OR, USA) (GenBank accession no. JN578697, target sequence: 5’ CATCACAGGTGGACAGCGGCATGGA 3’) in 0.2M KCl as described previously [29]. Embryos were cultured until swim-up stage (6 days post fertilization - dpf) in small recirculation system described in Cheng et al. (2020) at room temperature (21-22 °C) with UV water treatment. Fish were then transferred to plastic dishes and placed in incubator (22 °C). First feeding was initiated with baby brine shrimp.

Prior to germ cell transplantation, a two-year old male specimen of mirror carp (aquarium raised, BW 425 g) (Fig. 1A) was over-anaesthetized in MS222 (E10505), stunned and sacrificed by exsanguination. Body was thoroughly disinfected with 70% ethanol. Testes were carefully dissected after abdominal-lateral incision, weighted (30 g) and placed in phosphate-buffered saline (PBS). Small fragment of testis was fixed in Bouin’s solution for histological sectioning. Part of the testes (5g) was separated, cleaned off the big blood vessels and cut with scalpel to small (∼100 mg) fragments and rinsed in PBS several times to wash out leaking sperm. Afterwards, fragments were transferred into two 50ml tubes and cut finely with scissors. Tissue dissociation (25 ml of media was used per one 50ml tube) was done in media containing 0.15% trypsin (T4799) and 0.1% collagenase (C0130) diluted in PBS on a laboratory shaker for 1.5h at 20 °C. DNase I (10104159001) (aliquoted to 5% stock solution in RNase free water) was added continuously when clumping was observed, in total, 500 µl of DNase I was used. Tissue dissociation was terminated by adding 20ml of L15 media with 20% foetal bovine serum (FBS) per each 50ml tube and filtrated using 40 µm cell strainer (CLS431750-50EA). Due to the presence of large amount of sperm (Fig. 1B), 30% percoll gradient sorting was performed as described [16] to enrich testicular cell suspension (Fig. 1C). Small fraction of dissociated tissue was separated before the addition of L15 with FBS and was used for PKH26 staining as described previously [30]. Recipients receiving labelled cells were reared separately and were not reproduced.

**Figure 1.**
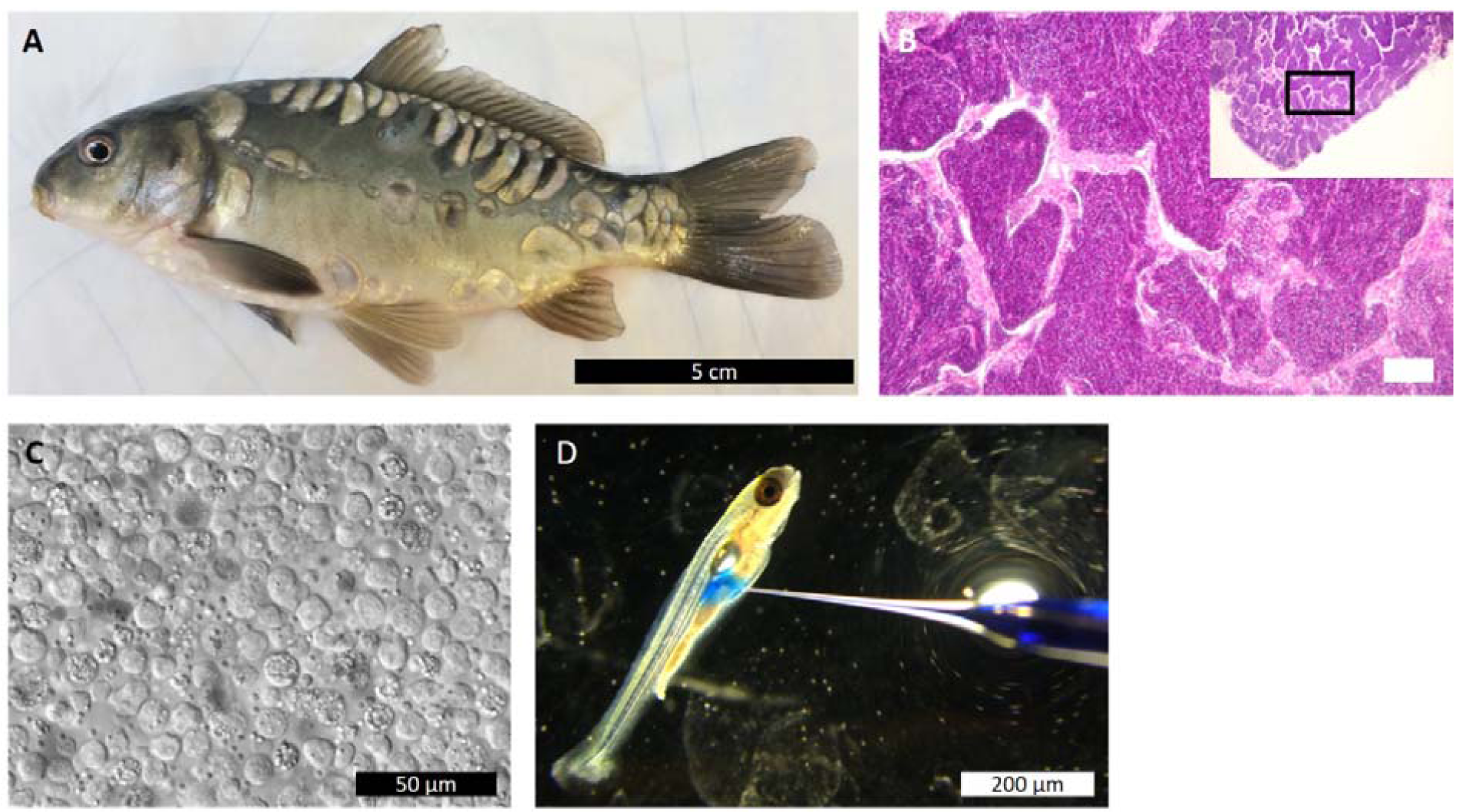
Cell transplantation procedure. A) Used male donor, B) Histological section from donor testes, predominantly containing spermatozoa. Caption in the left corner with black rectangle is low magnification of the presented view, scale bar 100 µm. C) Enriched testicular cell suspension after percoll gradient sorting. D) Intraperitoneal transplantation into goldfish (note that injected solution is only methylene blue solely for purpose of documentation).

Cell were transplanted into 7dpf MO treated goldfish, anaesthetized in 0.05% MS222 buffered with TRIS (TRIS-RO), and placed on agar coated Petri dish. Testicular cells suspension was loaded into glass microcapillaries which were previously opened using a capillary grinder (EG-401, Narishige, Japan). Capillary was attached to a micromanipulator and pneumatic injector (FemtoJet® 4i, Eppendorf, Germany). Each goldfish received 3-5 × 10^4^ testicular cells intraperitoneally when injection was positioned around the posterior part of the gas bladder (Fig. 1D).

### Germline chimeras rearing and reproduction

Transplanted goldfish larvae (n 100) were transferred to the aquarium and held constantly at 24-25 °C, 14L:10D photoperiod, fed by *Artemia* nauplii *ad libitum* and from 3 wpf co-fed with dry diet. Full transition to the dry diet was from 4wpf. At 4 months of age, goldfish were transferred into 200l aquaria. Fish were individually identified by PIT tags at age of 18 months and were transferred to bigger tank (500L) equipped with controlled heating/cooling and water filtration.

Reproduction of the goldfish germline chimeras was conducted at 2 years of age (2019) and repeated at 3 years of age (2020) together with common carp reproduction. For the sake of synchronization of gamete maturation, goldfish were exposed to temperature fluctuation. Temperature was decreased from initial 24 °C every week for 1-2 °C until 8 °C was reached and fish were held at this temperature for 6 weeks. After that, temperature was increased every week for 1-2 °C until 21 °C were reached. Photoperiod was managed accordingly by shortening light cycle down to 8 h during the cold phase and prolonging light cycle up to 14h at heating phase. Hormonal stimulation of all transplanted goldfish was conducted as it is described for goldfish females in two doses (0.5 and 2.5 mg of carp pituitary per kg of body weight). At the time of expected ovulation and spermiation all fish were checked by hand stripping. Sperm was collected using 1ml pipet with piston-type tips and stored on ice. Sperm volume was measured in a pipette and spermatozoa concentration was determined in Bürker chamber counting cells in 20 squares after dilution in Kurokura 180 solution for each chimeric male producing sperm and three control males of common carp and goldfish. Eggs were stripped into 50 ml tubes in case of transplanted goldfish and control goldfish females, carp eggs were firstly collected in plastic bowls and part of them was sampled in 50 ml tubes. Eggs were weighted and stored at 10 °C. PIT tags and weight of the fish producing gametes were recorded.

In 2019 two surrogate females produced a small amount of eggs. Randomly chosen sperm from five males was pooled and all eggs were fertilised. However, very low fertilisation rate was observed and we decided to fix surviving embryos for DNA analysis. Survival was not recorded in 2019. Sperm volume and spermatozoa concentration were recorded.

In 2020 eggs from transplanted goldfish females were obtained and used for fertilisation and genotyping by primers for RT-PCR. Five combination of fertilization were performed according to parental species in 2020 (Table 1). Small portions of collected eggs from surrogate females were fertilized individually with the individual sperm from goldfish transplanted males (n 5) and pooled common carp sperm. Egg/sperm ratio was 1g/5µl, gametes were activated with 0.5 ml of dechlorinated water and stirred for 20 seconds). Eggs were always fertilized on a petri dish (9 cm diameter), allowed to stick and then transferred to incubation system with controlled water temperature (21 °C) and held until hatching. Fertilization, eyeing and swim-up rate was recorded. Common carp eggs from Amur mirror carp strain were fertilised individually with sperm from all chimeric males. Pure common carp control was established as well. A representative sample of progeny from those five groups of crosses (Table 1) were pooled (100-200 individuals in each pooled group) at 14dpf and were reared in aquaria (200 l) at 23-24 °C for three months. Due to the limitation of incubation capacities, reproduction of control goldfish was performed two weeks after the chimera and carp reproduction. Pooled sperm from three goldfish male was used to fertilise eggs from three goldfish females individually. At that time, a small number of hybrids was produced by fertilising goldfish eggs with common carp sperm. Note that the performance of hybrid progeny was not monitored as they were used solely for phenotype documentation. Part of swim-up larvae was anesthetized and frozen fixed for later DNA extraction and genotyping. Eggs were fixed in 70% EtOH for calculation of relative and absolute fecundity (3 females per species, triplicated) and in 4% paraformaldehyde to measure egg diameter (3 females per species, triplicated). Part of the collected sperm from chimeric males 1,2 and 3, three goldfish males and three carp males was firstly diluted in Kurokura 180 [32] solution and then fixed in 2.5% glutaraldehyde in PBS for electron microscopy. Remainder of the sperm and eggs was frozen and stored for later analysis (DNA or RNA extraction).

**Table 1.** Parental combinations which were pooled after hatching and monitored for survival.

| Group | Sire | Dam |
| --- | --- | --- |
| GCxGC | Chimeric goldfish | Chimeric goldfish |
| CCxGC | Control carp | Chimeric goldfish |
| GCxCC | Chimeric goldfish | Control carp |
| CCxCC | Control carp | Control carp |
| GFxGF | Control goldfish | Control goldfish |

### Confirmation of donor-derived gamete and larvae origin

Total genomic DNA from sperm, larvae and fin clips was extracted by PureLink™ Genomic DNA Mini Kit (Invitrogen™) according to manufacturer’s instructions. Species-specific primers were designed according to NCBI using Primer-Blast. Selected primers were validated to be species-specific using fin clips from 5 specimens of common carp, goldfish and common carp (male) x goldfish (female) hybrids. Primers for DNA genotyping are listed in table 2. All sperm samples obtained from chimeric males (in both years), control carp and goldfish were genotyped. RNA from eggs of chimeric (n 8), carp (n 3) and goldfish (n 3) females reproduced in 2020 was extracted using PureLink RNA Mini Kit and treated with DNase (12185010, Thermo-Fisher). Isolated RNA was transcribed to cDNA using WizScript™ RT FDmix kit (Wizbiosolutions, South Korea). RT-PCR was done with carp and goldfish specific primers for *ddx4* (*vasa*) gene which is expressed in germ cells exclusively. Primers for *vasa* gene for RT-PCR with eggs are listed in table 3. The reaction mixture for PCR and RT-PCR contained 1 μl template DNA/cDNA, 0.5 μl forward and 0.5 μl reverse primer, 5 μl PPP Master Mix (Top-Bio) and 3 μl PCR H_2_O (Top-Bio). Reaction conditions were 30 cycles of 94 °C for 30 s, 58 °C for 30 s and 72 °C for 30 s. Products were analysed on 2% agarose gel using UV illuminator.

**Table 2.**
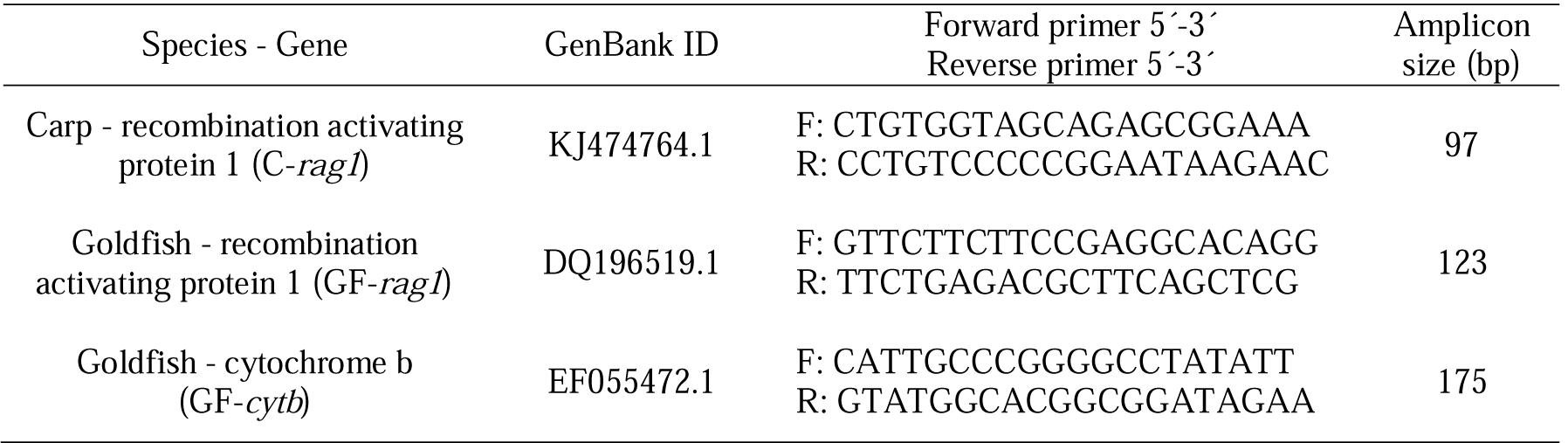
Species-specific primers for DNA genotyping.

**Table 3.**
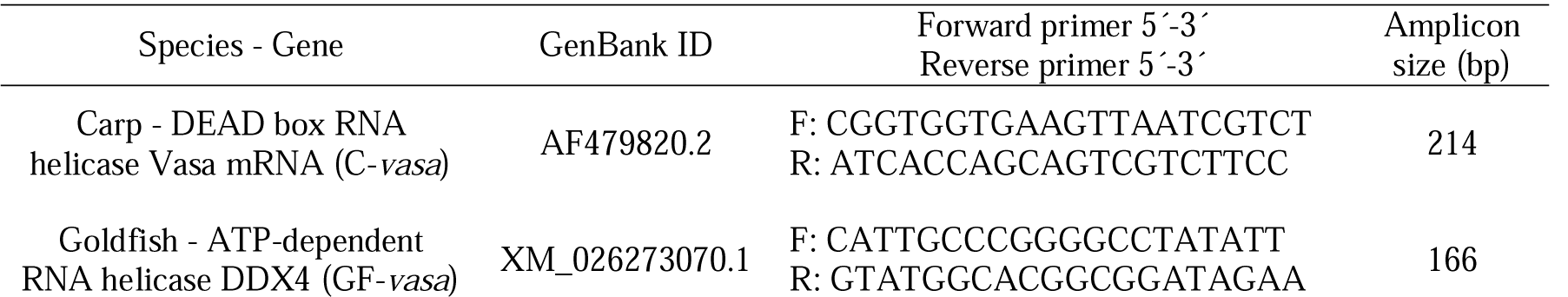
Species-specific primers for detection of mRNA in donor-derived and control oocytes.

### Histology

Testicular tissue from donor specimen was fixed overnight in Bouin’s fixative, washed in 70% ethanol, dehydrated and cleared in ethanol-xylene series, embedded into paraffin blocks, and cut into 5 mm thick sections. Mounted paraffin slides were stained with hematoxylin and eosin. Histological sections were photographed using a microscope (Nikon Eclipse Ci) with mounted camera (Canon EOS 1000D).

### Electron microscopy

Sperm samples for scanning electron microscopy (SEM) from three male chimeras (chimeric male 1, 2, and 3), three control goldfish and three control carps were firstly diluted in Kurokura 180 solution [32] and then primary fixed in 2.5% glutaraldehyde in PBS and stored in fridge until processing. The samples were stuck on poly-l-lysine-coated glass slides and secondary fixed in 4% osmium tetroxide for 2□h at 4□°C, washed three times in PBS, dehydrated gradually through acetone series (30, 50, 70, 90, 95 and 100% acetone, 15 min each), dried with a Pelco CPD2 CO_2_ Critical Point Dryer, mounted on a metal stub, coated with gold under vacuum with SEM coating unit E5100 (Polaron Equipment).

### Statistical analysis

Data with normal distribution were assessed using One-way ANOVA with Tukey’s honest significant difference. Data without normal distribution were assessed using Kruskal-Wallis ANOVA with Dunn’s multiple comparison test. Significance level was set for all trials at p < 0.05. Statistical analysis was performed using Statistica v13.1 software (TIBCO Inc., Palo Alto, CA, USA).

## Results

### Transplantation and germline chimera survival

After the transplantation, colonisation patterns of PKH 26 labelled cells were monitored occasionally. Day after the transplantation, a high number of fluorescent cells were visible alongside the gas bladder, mostly located at its caudal part (Fig. 2A). Noticeable decrease in number of fluorescent cells was observed one-week post-transplantation when only a few individual cells located in the close vicinity of the gas bladder (Fig. 2B). Dissection performed one-month post-transplantation showed fluorescent cells in proliferation phase according to their clustering in the presumptive gonads (Fig. 2C,D). Two weeks post-transplantation, 90 larvae were alive, at time of PIT tagging (18 mo) 72 fish were alive and all of them survived until the first reproduction in August 2019. Second reproduction in May 2020 was conducted with 71 surviving fish and all of them were alive at 3 months post reproduction.

**Figure 2.**
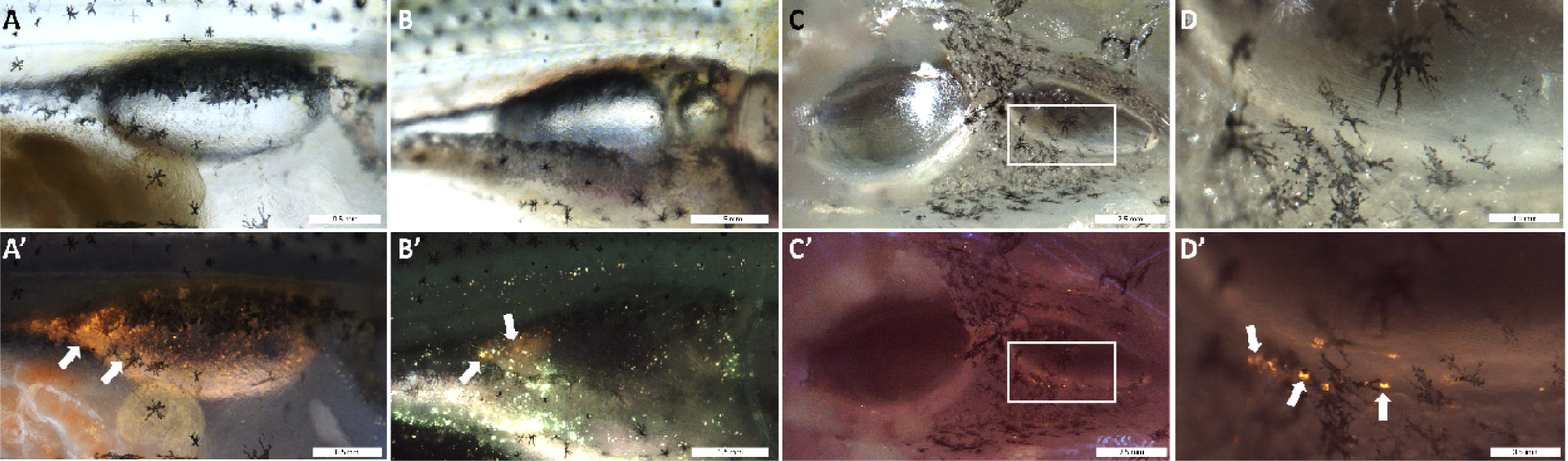
Colonization of the goldfish genital ridge by PKH26 labelled carp testicular cells. A) 24hpt, lateral view, B) 7dpt, lateral view, C and D) 30dpt, ventral view on dissected body cavity. Capital letters are brightfield captions, capital letters with apostrophe are captions of the same area taken using fluorescent filter DAPI/FITC/TRITC. Labelled cells have orange signal and are depicted with arrows. White rectangle in C and C’ depicting magnified view in D and D’.

### Germline chimera reproduction and genotyping

Sperm was successfully collected from 25 surrogate goldfish in various amounts and densities during the first reproduction in 2019. Genotyping with carp and goldfish specific primers revealed the presence of common carp DNA in all collected samples while goldfish specific amplicons were not detected in collected sperm. Two goldfish surrogate females ovulated low amount of eggs (chimeric female 1 ovulated 240 eggs and chimeric female 2 ovulated 360 eggs) later confirmed to be of donor-derived origin. Due to the low amount of the available eggs, pooled sperm from 5 males was used for individual fertilization in 2019. Unfortunately, all embryos showed delayed development and did not survive the first 24 hours. However, DNA from at least 10 embryos from each fertilization was isolated individually and PCR amplicons corresponded to pure common carp progeny in all genotyped embryos (n 123) meaning 100% sterilisation success. Remaining goldfish (n 48) did not produce any gametes.

During second reproduction of chimeras, gametes were obtained from 23 males and 8 females. Before gamete collection, courtship behaviour when chimeric males were chasing females was observed. Moreover, males developed typical tubercles on the operculum (Supplementary file 1). Remaining transplanted goldfish did not produce gametes and did not show sexual sings. Genotyping of obtained chimeric sperm with C-*rag1* primers showed presence of carp specific amplicons. Goldfish specific amplicons with GF-*cytb* primers were not detected in samples of chimeric and control carp sperm. Goldfish specific amplicons with GF-*rag1* primers were not detected in chimeric and control carp sperm, only one unspecific product with size about 650bp was detected, but they were not detected in goldfish controls (Fig. 3A). Later analysis of progeny from each cross between two chimeric parents or cross between chimeric parent and control carp showed presence of carp-specific amplicons only. Ten larvae were genotyped from each parental combination where germline chimera was included (note that only one larva survived in combination of chimeric female 2 with chimeric male 5). Thus, 421 larvae were genotyped in total. Primers for GF-*cytb* were not used for progeny genotyping because potential contribution of paternal mitochondrial DNA would not be detected in the progeny. Carp x goldfish hybrids could be detected by presence of both amplicons for C-*rag1* and GF-*rag1* primers (Fig. 3B). RT-PCR with eggs from all chimeric females producing eggs (n 8) showed presence of carp *vasa* amplicon only (Fig. 3C). Thus, we can conclude, that only donor-derived gametes were obtained from all chimeras including that five low-egg producing chimeric females. In result, pure donor-derived common carp progeny was obtained.

**Figure 3.**
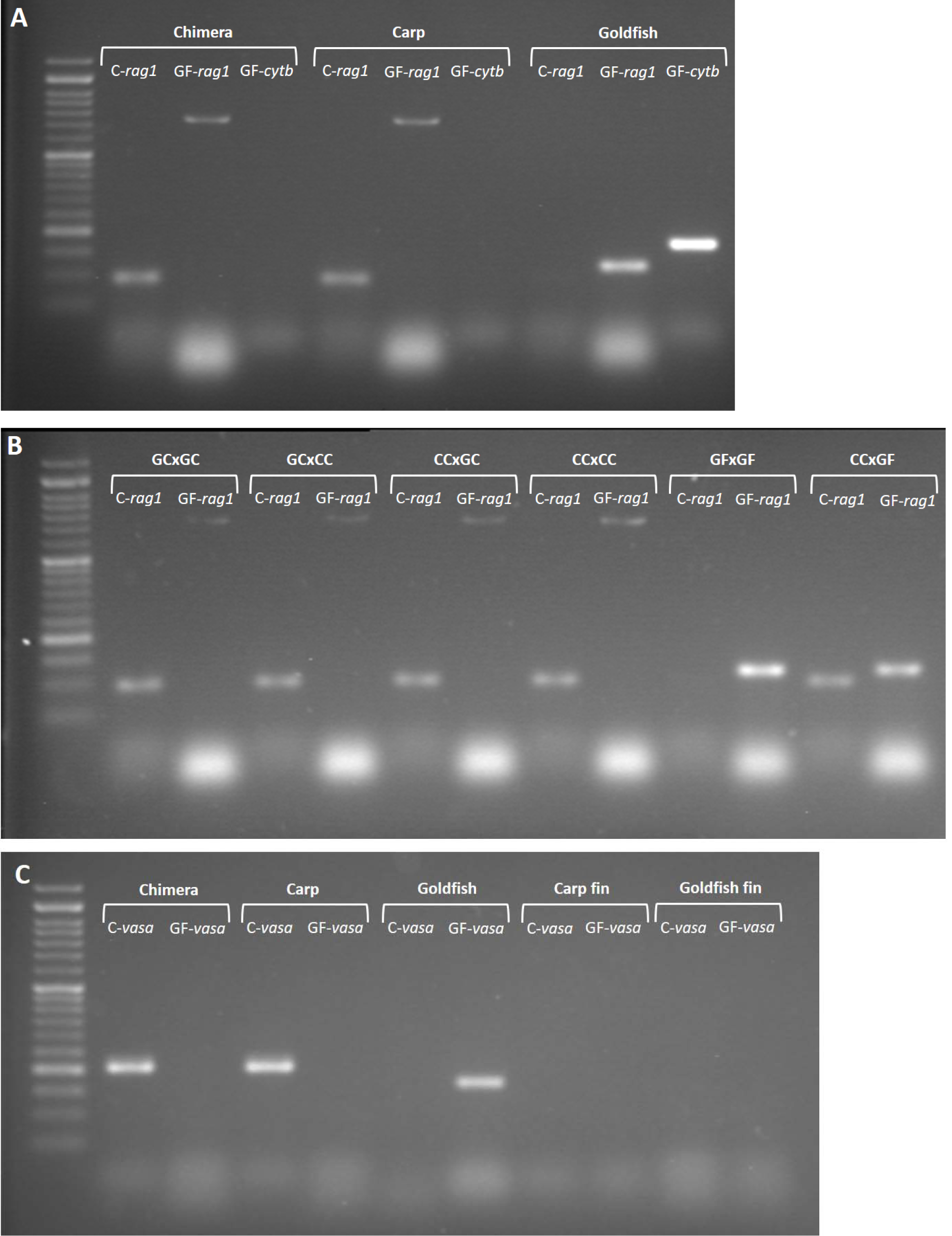
Detection of donor-derived gamete production and offspring identification by species-specific primers. A) Gel electrophoresis of PCR products from DNA obtained from sperm of chimera, carp and goldfish. B) Gel electrophoresis of PCR products from DNA obtained from larvae of different crosses parental crosses. Abbreviations above white brackets stand for parental combination: GC – germline chimera, CC – common carp, GF – goldfish. C) Gel electrophoresis of RT-PCR products from eggs obtained from chimera, carp and goldfish female with fin tissue controls.

### Reproductive characteristics

Individual goldfish female chimeras produced significantly lower number of eggs in gram compared to carp but higher number compared to goldfish control females (Fig. 4A). Relative fecundity calculated as the number of eggs per gram of body weight was not significantly different between chimeras and control goldfish but was lower from relative fecundity of control carp females (note that species were statistically compared, not the individuals). Same pattern was observed when absolute fecundity was compared (Tab. 4). The diameter of unfertilised eggs of chimeras was significantly different from carp control eggs comparing the species (Fig. 4B). The diameter of fertilised chimeric eggs was significantly different from both control species when the size of chimeric eggs was between those from carp and goldfish. However, significant differences were also found comparing individual females (Fig. 4C). Egg characteristics of chimeric females can be considered to be between eggs of common carp and goldfish. Regarding the colour, chimeric eggs under the natural light appeared to be from yellowish to greenish, corresponding to the normal range of colours observed in common carp and goldfish (Fig. 5A-C).

**Figure 4.**
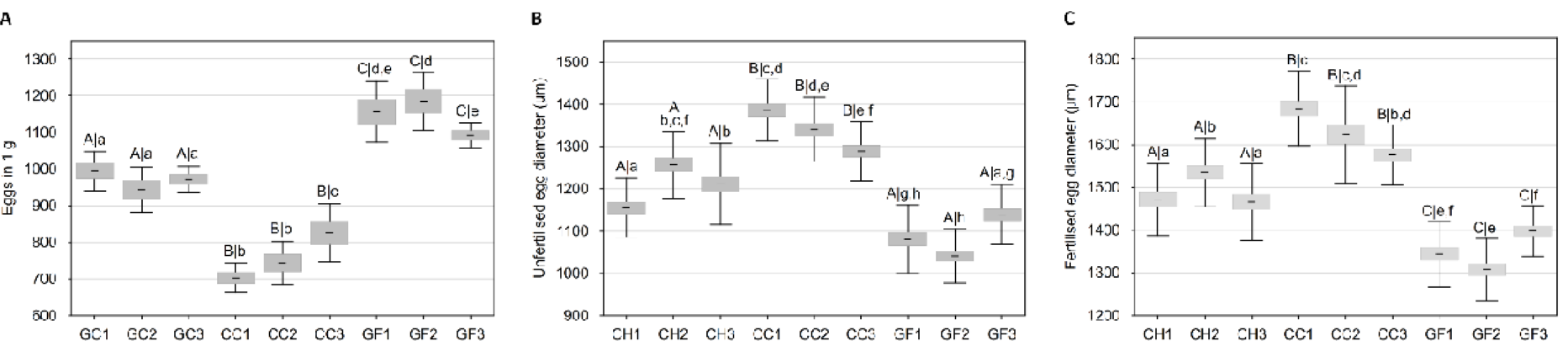
Egg characteristics in females of germline chimeras (GC1-3), common carps (CC1-3) and goldfish (GF1-3) controls. A) Amount of dry egg in one gram; B) Diameter of unfertilised eggs (fixed in 4% PFA); C) Diameter of fertilised eggs (4-5 hours post fertilisation). Data are presented as a mean ± confidence interval with SD. Different capital letters stand for differences in species, small letters stand for individual differences.

**Figure 5.**
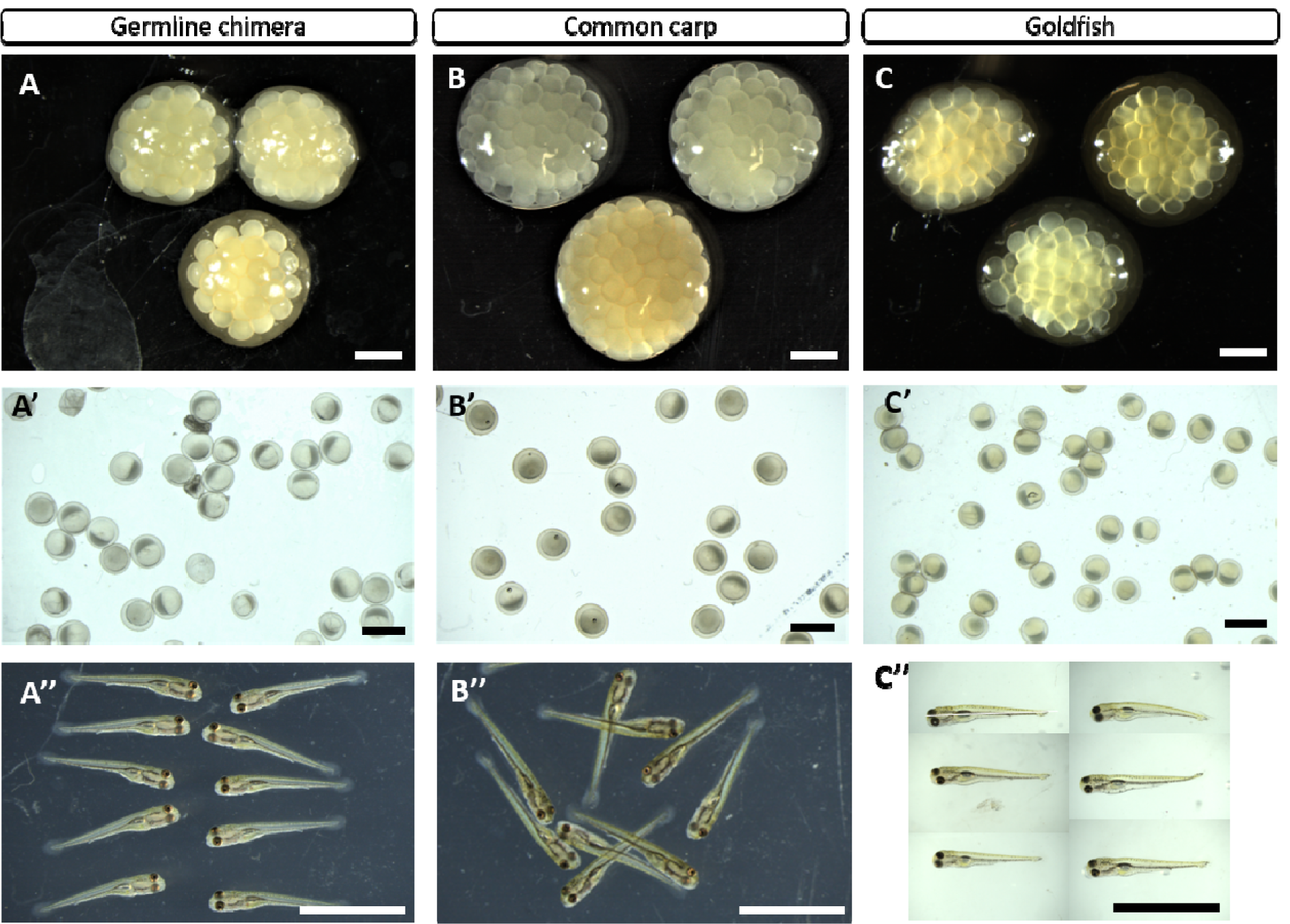
Gametes and progeny from chimeras and controls. A-C) Representative portion of oocytes obtained from chimeric (A), carp (B) and goldfish females (C). Representative captions of chimeric (A’), carp (B’) and goldfish embryos (C’) at 4-5 hpf. Swim-up progeny of germline chimeras (A’’), carp (B’’) and goldfish (C’’). Scale bars A-B and A’-C’ 2 mm, A’’-C’’ 1 mm.

Sperm was collected from 25 fish in 2019 and from 23 fish in 2020 (note that one chimeric male deceased 3 months after the first reproduction in August 2019 and one male which spermiated in 2019 did not spermiate in 2020). Chimeric males produced sperm in various volumes and various concentration of spermatozoa. Compared to goldfish control males, sperm concentration was about 1/3 lower and about half when compared to control carp males. Similar pattern was observed in the total volume of collected sperm from chimeras which was about 1/3 lower compared goldfish control males and ∼50 times lower compared to carp control males. Complete data on male reproductive performance over two years are given in table 5. Width and length of the spermatozoa head of germline chimeras showed some variability across individual males, however, species did not differ significantly (Fig. 6A,B). Distinct characteristics were observed in flagellum length of spermatozoa from chimeric males and control carps when both were significantly shorter to spermatozoa of goldfish control males after comparing individually and by species (Fig. 6C). During observation on SEM, no abnormalities such as biflagellate spermatozoa or abnormally shaped heads were observed in any of specimens. Representative spermatozoa captions are given in figure 7.

**Table 4.**
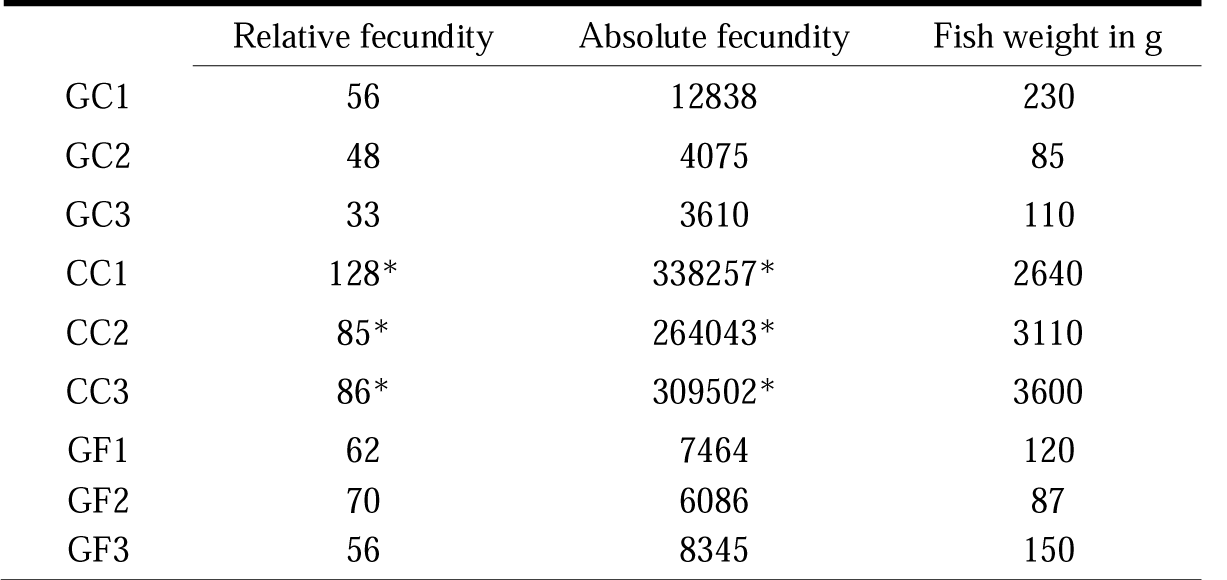
Relative and absolute fecundity of germline chimera (GC1-3), common carp (CC1-3) and goldfish (GF1-3) females during reproduction in 2020. Asterisks stand for significant difference in species.

|  | Relative fecundity | Absolute fecundity | Fish weight in g |
| --- | --- | --- | --- |
| GC1 | 56 | 12838 | 230 |
| GC2 | 48 | 4075 | 85 |
| GC3 | 33 | 3610 | 110 |
| CC1 | 128* | 338257* | 2640 |
| CC2 | 85* | 264043* | 3110 |
| CC3 | 86* | 309502* | 3600 |
| GF1 | 62 | 7464 | 120 |
| GF2 | 70 | 6086 | 87 |
| GF3 | 56 | 8345 | 150 |

**Table 5.**
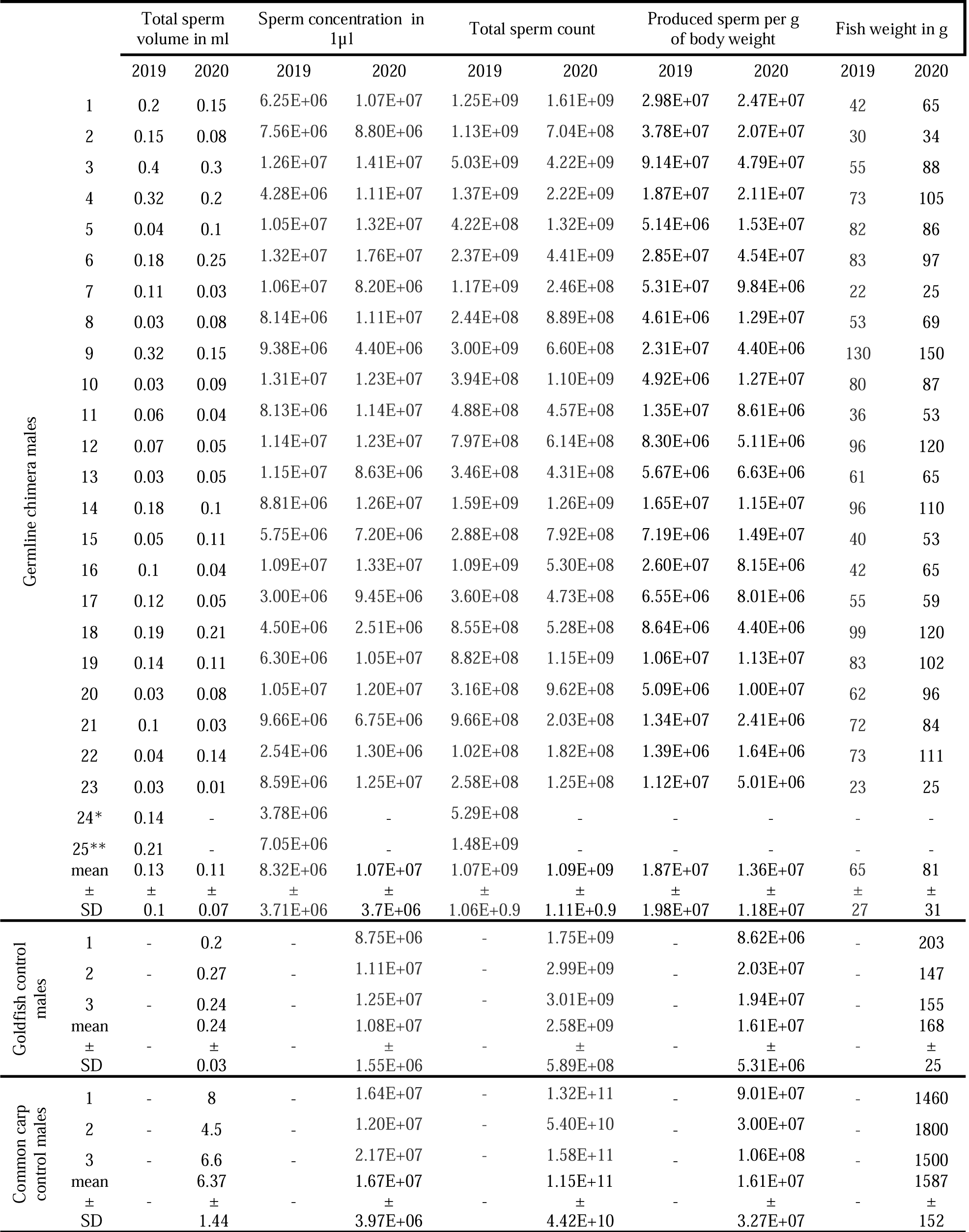
Reproductive characteristics of males during two reproduction in 2019 and 2020. * male 24 did not spermiate in 2020. ** male 25 died between spawning in 2019 and 2020.

**Table 6.** Reproductive performance of various parental combinations and common carp and goldfish controls during second reproduction in 2019.

| Female | Male | Eggs | Fertilization rate |  | Eyeing rate |  | Swim-up rate |  |
| --- | --- | --- | --- | --- | --- | --- | --- | --- |
|  |  |  | n | % | n | % | n | % |
| Chimera female 1 | Male chim. 1 | 321 | 307 | 95.6 | 296 | 92.2 | 170 | 53.0 |
|  | Male chim. 2 | 280 | 267 | 95.4 | 257 | 91.8 | 154 | 55.0 |
|  | Male chim. 3 | 242 | 239 | 98.8 | 237 | 97.9 | 184 | 76.0 |
|  | Male chim. 4 | 276 | 249 | 90.2 | 244 | 88.4 | 180 | 65.2 |
|  | Male chim. 5 | 249 | 223 | 89.6 | 213 | 85.5 | 176 | 70.7 |
| | Summary MEAN ± SD | | 93.9 ± 3.5 | | 91.2 ± 4.2 | | 64 ± 8.9 | |
|  | Carp males mix | 260 | 255.0 | 98.1 | 230.0 | 88.5 | 180.0 | 69.2 |
| Chimera female 2 | Male chim. 1 | 154 | 139 | 90.3 | 114 | 74.0 | 12 | 7.8 |
|  | Male chim. 2 | 258 | 169 | 65.5 | 164 | 63.6 | 71 | 27.5 |
|  | Male chim. 3 | 201 | 148 | 73.6 | 146 | 72.6 | 58 | 28.9 |
|  | Male chim. 4 | 186 | 123 | 66.1 | 119 | 64.0 | 38 | 20.4 |
|  | Male chim. 5 | 124 | 120 | 96.8 | 120 | 96.8 | 1 | 0.8 |
| | Summary MEAN ± SD | | 78.5 ± 12.8 | | 74.2 ± 12.1 | | 17.1 ± 11 | |
|  | Carp males mix | 105 | 69.0 | 65.7 | 42.0 | 40.0 | 18.0 | 17.1 |
| Chimera female 3 | Male chim. 1 | 219 | 201 | 91.8 | 191 | 87.2 | 157 | 71.7 |
|  | Male chim. 2 | 129 | 128 | 99.2 | 125 | 96.9 | 82 | 63.6 |
|  | Male chim. 3 | 126 | 125 | 99.2 | 123 | 97.6 | 63 | 50.0 |
|  | Male chim. 4 | 323 | 302 | 93.5 | 302 | 93.5 | 228 | 70.6 |
|  | Male chim. 5 | 104 | 104 | 100.0 | 100 | 96.2 | 79 | 76.0 |
| | Summary MEAN ± SD | | 96.7 ± 3.4 | | 94.3 ± 3.8 | | 66.4 ± 9.1 | |
|  | Carp males mix | 147 | 142.0 | 96.6 | 133.0 | 90.5 | 82.0 | 55.8 |
| Carp female 1 | Male chim. 1 | 209 | 207 | 99.0 | 205 | 98.1 | 187 | 89.5 |
|  | Male chim. 2 | 214 | 213 | 99.5 | 211 | 98.6 | 200 | 93.5 |
|  | Male chim. 3 | 229 | 228 | 99.6 | 224 | 97.8 | 203 | 88.6 |
|  | Male chim. 4 | 257 | 249 | 96.9 | 243 | 94.6 | 183 | 71.2 |
|  | Male chim. 5 | 274 | 274 | 100.0 | 272 | 99.3 | 216 | 78.8 |
| | Summary MEAN ± SD | | 99 ± 1.1 | | 97.7 ± 1.6 | | 84.3 ± 8.1 | |
|  | Carp males mix | 303 | 290 | 95.7 | 273 | 90.1 | 249 | 82.2 |
| Carp female 2 | Male chim. 1 | 168 | 168 | 100.0 | 168 | 100.0 | 144 | 85.7 |
|  | Male chim. 2 | 266 | 265 | 99.6 | 242 | 91.0 | 232 | 87.2 |
|  | Male chim. 3 | 196 | 196 | 100.0 | 192 | 98.0 | 178 | 90.8 |
|  | Male chim. 4 | 205 | 192 | 93.7 | 192 | 93.7 | 157 | 76.6 |
|  | Male chim. 5 | 305 | 303 | 99.3 | 300 | 98.4 | 240 | 78.7 |
| | Summary MEAN ± SD | | 98.5 ± 2.4 | | 96.2 ± 3.3 | | 83.8 ± 5.3 | |
|  | Carp males mix | 289 | 280.0 | 96.9 | 273.0 | 94.5 | 248.0 | 85.8 |
| Carp female 1 | Male chim. 6 | 172 | 172 | 100.0 | 172 | 100.0 | 163 | 94.8 |
|  | Male chim. 7 | 178 | 177 | 99.4 | 176 | 98.9 | 156 | 87.6 |
|  | Male chim. 8 | 152 | 151 | 99.3 | 148 | 97.4 | 145 | 95.4 |
|  | Male chim. 9 | 186 | 185 | 99.5 | 185 | 99.5 | 172 | 92.5 |
|  | Male chim. 10 | 215 | 214 | 99.5 | 210 | 97.7 | 208 | 96.7 |
|  | Male chim. 11 | 221 | 208 | 94.1 | 197 | 89.1 | 190 | 86.0 |
|  | Male chim. 12 | 247 | 235 | 95.1 | 227 | 91.9 | 195 | 78.9 |
|  | Male chim. 13 | 206 | 181 | 87.9 | 177 | 85.9 | 156 | 75.7 |
|  | Male chim. 14 | 225 | 210 | 93.3 | 201 | 89.3 | 186 | 82.7 |
|  | Male chim. 15 | 199 | 184 | 92.5 | 182 | 91.5 | 162 | 81.4 |
|  | Male chim. 16 | 215 | 198 | 92.1 | 198 | 92.1 | 179 | 83.3 |
|  | Male chim. 17 | 256 | 241 | 94.1 | 237 | 92.6 | 213 | 83.2 |
|  | Male chim. 18 | 191 | 179 | 93.7 | 177 | 92.7 | 159 | 83.2 |
|  | Male chim. 19 | 196 | 186 | 94.9 | 186 | 94.9 | 169 | 86.2 |
|  | Male chim. 20 | 218 | 192 | 88.1 | 188 | 86.2 | 164 | 75.2 |
|  | Male chim. 21 | 143 | 132 | 92.3 | 130 | 90.9 | 127 | 88.8 |
|  | Male chim. 22 | 249 | 230 | 92.4 | 227 | 91.2 | 191 | 76.7 |
|  | Male chim. 23 | 172 | 170 | 98.8 | 165 | 95.9 | 138 | 80.2 |
| | Summary MEAN ± SD | | 94.8 ± 3.7 | | 93.2 ± 4.2 | | 84.9 ± 6.5 | |
| Goldfish female 1 | Goldfish males mix | 248 | 235 | 94.8 | 221 | 89.1 | 204 | 82.3 |
| Goldfish female 2 |  | 196 | 162 | 82.7 | 151 | 77.0 | 145 | 74.0 |
| Goldfish female 3 |  | 301 | 282 | 93.7 | 257 | 85.4 | 228 | 75.7 |
| | Summary MEAN ± SD | | 90.4 ± 5.5 | | 83.8 ± 5.0 | | 77.3 ± 3.6 | |

**Figure 6.**
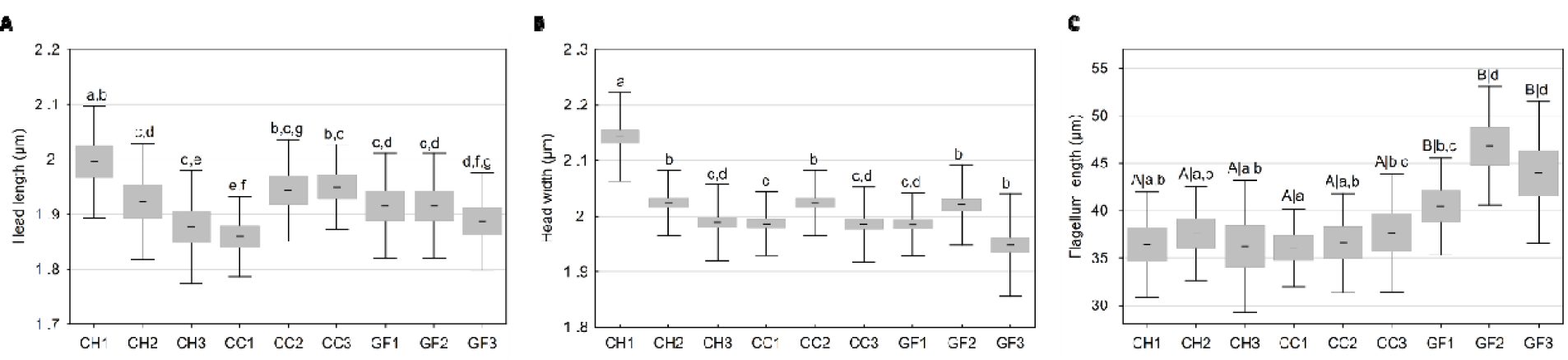
Spermatozoa morphological characteristics in males of germline chimeras (GC1-3), common carps (CC1-3) and goldfish (GF1-3) controls. A) Head length; B) Head width; C) Flagellum length. Data are presented as a mean ± confidence interval with SD. Different capital letters stand for differences in species, small letters stand for individual differences.

**Figure 7.**
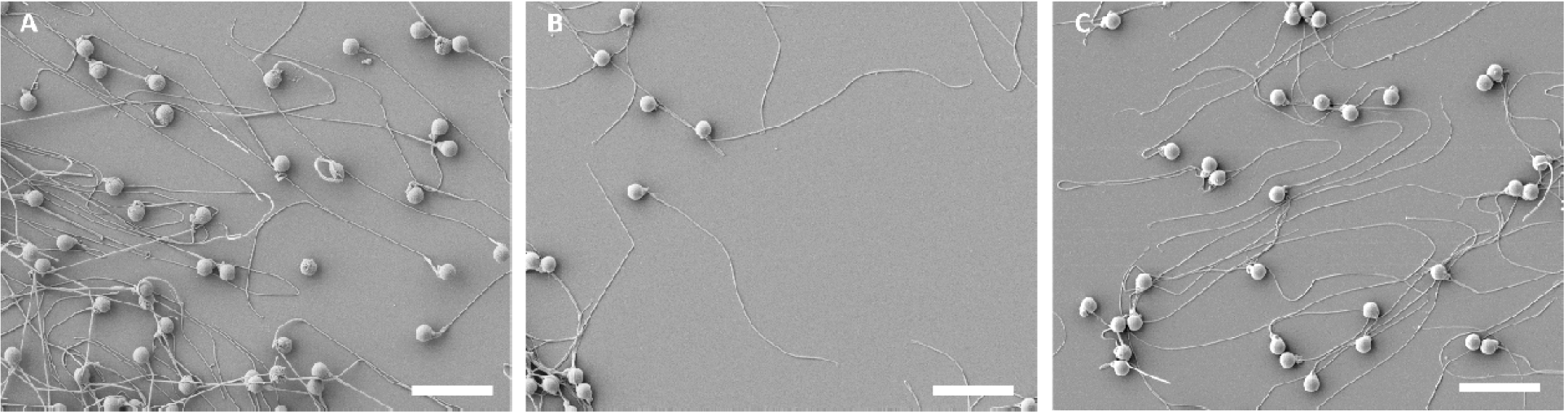
SEM images of spermatozoa from males A) Germline chimera male 1; B) Common carp male 1; C) Goldfish male 1. Scale bars 10 µm.

Fertilisation rates were comparable between two chimeric females regardless of the sperm origin (from chimeric males or carp males). However, female chimera 2 exhibited lower fertilisation rate and later on very low swim-up rate. Comparing species, summarized mean values of the embryo survival in chimeric females were slightly lower compared to the carp and goldfish controls (Tab. 6). Embryos from chimeras hatched 3-4 days post fertilization (dpf), swim-up stage was reached 5-6 dpf, same patterns were observed in control common carp and goldfish embryos. The external appearance of donor-derived larvae was similar to control common carp (Fig. 5A’’-C’’). After feeding initiation, only sporadic mortality occurred in the pooled groups. At 12 wpf, survival in all five pooled groups was over 95%. At one-month post fertilisation, scale patters could be identified by the naked eye when pure chimeric progeny, combination of one chimeric parent and control carp parent and common carp control exhibited mirror phenotype only (Fig. 8A-D). Majority of the body was nude and only a few bigger scales were observed. Contrary, control of goldfish (Fig. 8E) and common carp x goldfish hybrids (Fig. 8F) were distinguishable by scaly phenotype when body was covered with smaller scales entirely. Another distinct characteristic was in the shape of the dorsal fin. Carps had concave shape regardless their parents (Fig. 8A’-D’), goldfish had a straight shape (Fig. 8E’) and dorsal fin of the hybrid was very slightly concave (Fig. 8F’). Edge of the goldfish anal fin was perpendicular to the frontal plane of the body (Fig. 8E’’) while carp and hybrid had concave or convex shape but not perpendicular. Caudal fin in common carps was more forked than in the goldfish (Fig. 8E’’). However, we consider scale patterns and shape of the dorsal fin as a most distinctive feature for the first weeks. At 8 wpf, presence of barbells in common carps could be observed under a stereomicroscope, while goldfish and hybrid controls did not develop barbells.

**Figure 8.**
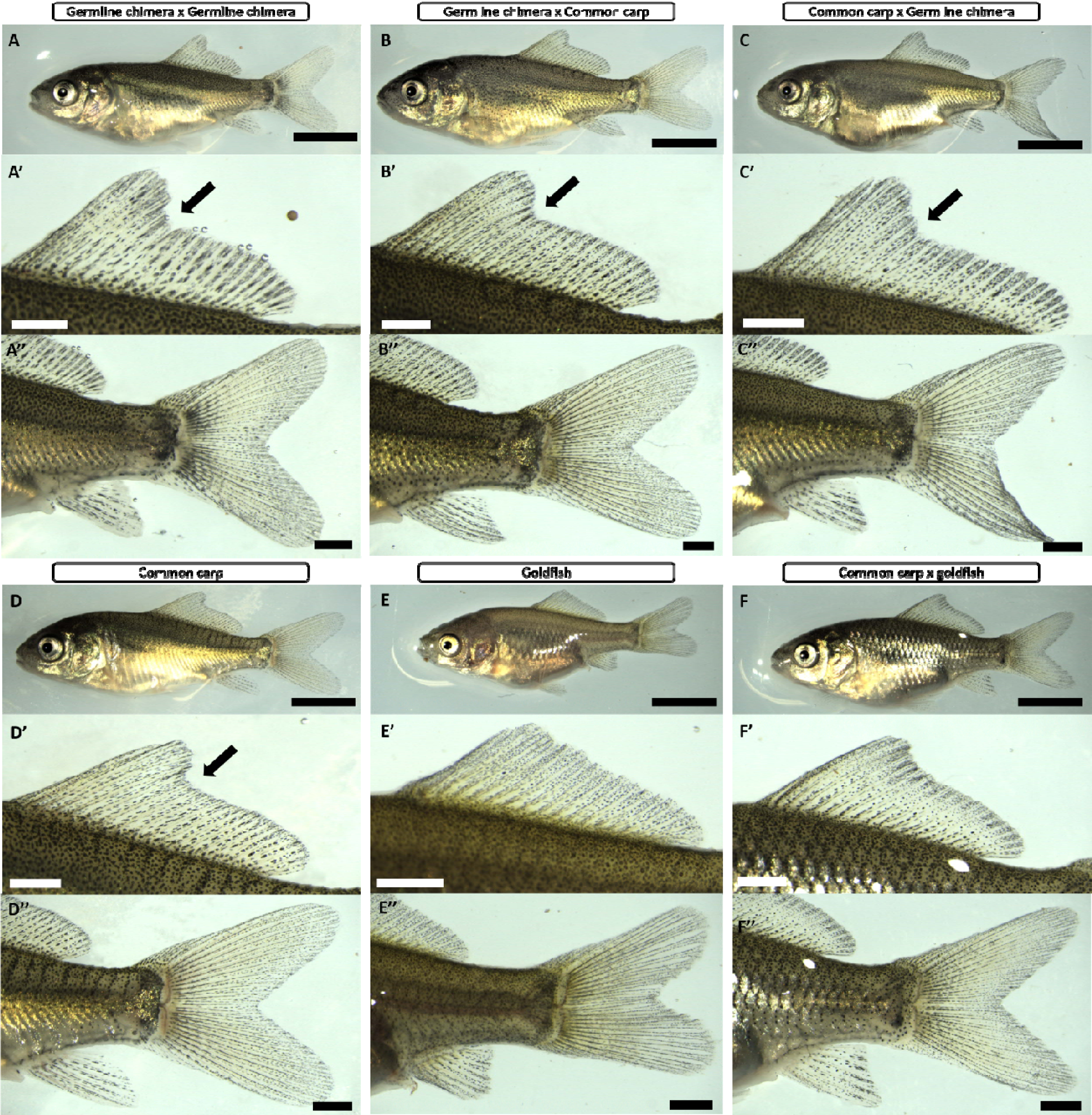
Appearance of the chimeric progeny with common carp and goldfish controls. A) Progeny from donor derived eggs and sperm; B) Progeny from cross between chimeric female and carp male; C) Progeny from carp female and chimeric male; D) Common carp control; E) Goldfish control; F) Hybrid progeny from cross between carp male and goldfish female. Letters A’- F’ are close captions of dorsal fin with arrows indicating concave shape typical for common carp, while goldfish has a straight shape and hybrid has less concave shape compared to the common carp. Letters A’’- F’’ are close captions of the caudal region. Scale bars A-F 4mm. A’-F’ and A’’-F’’ 1 mm.

## Discussion

Viable donor-derived common carp progeny from surrogate goldfish recipients transplanted by testicular germ cells was produced in this study. Moreover, we confirmed, that surrogate goldfish are capable to reproduce repeatedly when sperm and eggs were obtained in two consecutive years. From one hundred transplanted goldfish larvae, 33 of them produced donor-derived gametes when most of the surrogates matured at 2 years of age and reproduced again at 3 years of age which corresponds with the reproductive cycle of common carp and goldfish maturing in 2-4 year of age depending on conditions. The use of surrogate goldfish conveniently overcome the issue of big body sized broodstock of common carp which might be difficult to be kept in high numbers in fully controlled conditions. Largest chimeric female weighed 230g and smallest chimeric male producing sperm weighted 21 grams resulting in approximately 10-100 times smaller body size compared to common carp broodstock weighting 1.5-4 kg at the time of their first reproduction in our conditions (Supplementary file 2). This gives a possibility to conduct ongrowing and reproduction in reasonable space while allowing full control of the fish. The efficiency and suitability of this method is based on the fact that GSCs only from one donor are sufficient to establish production of viable progeny. We also need to point out, that size of the donor gonads would be sufficient to transplant into more than 500 of goldfish larvae, suggesting a large number of potential surrogates.

High numbers donor-derived sperm producing surrogates represent a solid backup of genetic resources from a putatively very precious single donor as they can be kept in different places. Moreover, surrogacy based on transplantation of stem cells of superior genotypes identified in selective breeding programs allows utilization of these superior genotypes in next generations of breeding schemes. Another yet undiscovered application of surrogacy is improved disease resistance. Currently, carp stocks are under threat of highly contagious koi herpes virus [33]. Virus can cause significant loses in market fish, but also to broodstock. According to literature, goldfish is not susceptible to koi herpes virus, moreover, presence of the virus was not detected in tissues of two goldfish strains [34]. Therefore, there is a great potential, that common carp germ cells can be protected in vivo using goldfish surrogates.

We can conclude, that previously reported 40-60% transplantation success of testicular [29] and ovarian [30] GSC in common carp is corresponding with numbers of adult germline chimeras producing donor-derived progeny in our study. Therefore, we can retrospectively say, that transplantation success assessed 2-3 months post transplantation probably gives reliable numbers about expected numbers of adult chimeras. Regarding the sex ratio, both species in this study are male heterogametic. Goldfish has been proven to have germ cell independent sex determination [35], but with the evident influence of higher temperatures during first weeks of life causing male bias. In our study, 75% of the reproduced chimeras were males which can be caused by relatively higher temperature during the ongrowing (24-25 °C). We also cannot exclude possibility that transplanted testicular cells might tend to follow their testicular fate even in the body of xenogeneic recipient. Also, potential role of other testicular somatic cells (Sertoli and Leydig) co-transplanted within cell suspension is not known yet and should be subject of further investigations. It is also realistic to expect that more chimeric females will appear because it seems that additional time is necessary for their maturation compared to chimeric males.

Intraspecific transplantation is challenging with respect to production of oocytes in good quality and sufficient amount [11,36] especially in more distant species [17]. Evolutionary distance between goldfish and carp is about 34 mya. Compared to the most recent summarization of surrogacy in fish by Goto and Saito [11], we obtained viable donor-derived progeny from most distant species so far. The greatest phylogenetic divergence in term of successful production of donor-derived eggs has been achieved between Atlantic salmon and rainbow trout which diverged 46 mya but obtained eggs did not give rise to viable progeny [17]. Two goldfish chimeric females produced thousands of oocytes with fertilization rates over 70% which could be regarded as very acceptable. Moreover, reproductive output of the large chimeric female 1 is promising from the point of the absolute fecundity which is hand to hand with increasing body weight. Thus, we intend to further monitor reproductive performance of surrogates to confirm their long-term suitability for genetic resources preservation. Also, we do not neglect those females producing a small amount of eggs. We suspect, that those low egg producing females might mature later and follow the patterns of two females producing a small amount of egg in 2019, but large amount of egg in 2020.

Assessed gamete characteristics from goldfish surrogates provided interesting findings on how host’s gonadal environment might influence gametogenesis. Most importantly, we found, that donor-derived spermatozoa have carp-like flagellum. This can be attributed to flagellum genesis, which is regulated autonomously on a single-cell level. Assembly of the flagellum is based on elements synthesis in the cell. Elements are then transported through the flagellar compartment towards to flagellar tip causing its elongation [37]. Next steps should be taken to asses physiology of donor-derived spermatozoa to observe if there is any causality of host gonadal environment on sperm motility parameters.

The eggs characteristics are more influenced by the host’s environment. This can be attributed to the fact, that oogenesis is compared to spermatogenesis more complex process requiring cell growth and transport of maternal deposits inside the oocytes. It is known that fish egg size, in general, can be very variable due to female-specific, but also environmental factors [38]. There is little information on how interspecific surrogacy can affect egg characteristics. Saito et al. [39] clearly showed that donor-derived oocytes are similar to the oocytes of the host species in terms of size. In our study, the size of donor-derived oocytes was between donor common carp and host goldfish oocytes. This can suggest certain influence of the host’s ovarian environment, but also genetic predetermination given on a level of single spermatogonia which differentiated into eggs. It will be very informative to provide a thorough assessment of egg nutrients compositions. We also need to be aware, that donor-derived oocytes are likely to contain deposits of maternal biomolecules.

The question of shortening the life cycle by surrogates was not the aim of this study, because both species have very similar reproductive biology. However, we need to point out, that this question on shortened life cycle via surrogates has not been answered in fish completely, because species with significant differences in maturity must be used. From our experience, carps or goldfish in temperate condition might start to spermiate spontaneously before reaching the first year, meaning, that intensive culture condition can rapidly accelerate the onset of maturity. The only robust answer to the potential of breeding acceleration could be probably given by surrogacy between extremely long maturing sturgeon species such as beluga transplanted into short maturing sterlet (difference in maturation is 15-20 years). Or utilise turquoise killifish (*Nothobranchius furzeri*) maturing in less than 3 weeks post hatching [40] as a recipient for longer maturing species.

Nowadays, wide pallet of GSCs transplantation methods as well sterilization methods became available and verified on several fish species [11,41]. Our strategy is based on complete depletion of endogenous PGCs via *dead end* gene knockdown as this gene is essential for PGCs migration and maintenance and its suitability for sterilization has been documented in several fish species [42]. It is necessary to mention, that the delivery method by microinjection is time demanding. However, we confirmed, that PGC depletion by MO is efficient and initial time and labour demands worth for complete sterilization success. We showed complete sterilisation by genotyping of gametes, embryos and fish and also by phenotypical characteristics when no progeny from surrogate parents showed full scale patterns typical for hybrids or goldfish. Other options for sterilization with subsequent transplantation in cyprinids such as triploidy induction has been tested, but in distant species from carp or goldfish such as zebrafish [43]. Data about gonadal development in triploid goldfish are not known but we presume, that gonads might have similar development as it was described in triploid koi variety of common carp capable to produce aneuploid gametes in large quantities [44] probably making it not suitable for surrogacy. Inconvenience of the triploid as recipient is known in other fish species. Triploid grass puffers (*Takifugu niphobles*) surrogates produced a mixture of donor-derived, but also own gametes after GSCs transplantation [45] which complicates direct application of surrogacy in aquaculture. Therefore, strategies targeting *dnd* gene are likely to facilitate sterilisation in wide range of species. On the other hand it is necessary to mention, that genomic resources are necessary to target *dnd* gene essential for PGCs migration by knock down or knock out techniques [42]. Hybridisation of common carp or goldfish might be another potential tool as hybrids were used successfully as surrogates in different species [46,47]. Performed studies reported sterility in cross of common carp a crucian carp [48] but no data are available regarding sex ratio of sterile hybrids. Sterile all male population of hybrids was produced by crossing goldfish females with common carp supermales (YY) [49], however, utilisations of this hybrid will probably lead to production of male germline chimeras only. Other numerous combinations of cyprinid hybrids were attempted, but often resulting in high mortality or abnormality rates accompanied with lack of data regarding sex ratio and gonadal development [50].

Purpose of the study was to provide an alternative approach for the production of donor-derived gametes of important aquaculture species. However, presented findings could be relevant for species conservation when surrogacy can be used to propagate threatened species [14,51,52]. A close relative of goldfish – crucian carp (*Carassius carassius*) is facing threats of population decline because of habitat loss and concurrence from its close but invasive relative – Prussian carp (*Carassius gibelio*) [53]. Efforts will be taken to develop strategies for preservation of crucian carp germ cells and their recovery in surrogate goldfish. Suitability of goldfish recipient can be suggested and justified by the evolutionary distance between crucian carp and goldfish (17 mya) which is approximately half of the distance between carp and goldfish (34 mya).

Reproductive performance of goldfish germline chimeras in this study gives a promising prospect for further analysis about the long-term reproductive performance of surrogates. More advanced application of surrogacy such as production of donor-derived gametes from cryopreserved germ cells, or production of monosex female stocks based on oogonia transplantation carrying XX chromosomes will be attempted further. Of the utmost importance, results of this study give us a positive outlook for oncoming reproduction of goldfish chimeras which received cells from a homozygous individual when production of genetically identical gametes and creation of an isogenic line is foreseen [54]. On the other hand, potential of highly fertile common carp recipients will be advantageous to rapidly increase production of small goldfish.

## Conclusions

Successful and repeated production of donor-derived common carp gametes from many times smaller goldfish surrogates clearly demonstrated convenience of the surrogate reproduction in one of the most important aquaculture species worldwide. Repeated reproduction of goldfish surrogates indicated the stemness properties and sexual plasticity of intraperitoneally transplanted carp testicular cells, when both sperm and eggs were obtained. Fertility performance of surrogates was close to goldfish controls, suggesting that introduced GSCs are capable to recover gametogenesis efficiently in PGCs depleted host. Presented technology in this study is ready to ameliorate preservation and breeding of common carp.

## Acknowledgement

Authors would like to thank to Michaela Fučíková, Christoph Steinbach and to staff of the Genetic Fisheries Centre for support and assistance during experimental work.

## Authors’ contributions

RF and MP conceived and designed the experiments; RF, VK and MP mainly performed the experiments; RF analysed the data; VK, DG and MP contributed with fish, reagents and materials; RF wrote the manuscript. All authors revised and approved the final version of the manuscript. RF had primary responsibility for final content of the manuscript.

## Availability of data and materials

The datasets used and/or analysed during the current study are available from the corresponding author on request.

## Ethics approval

Experimental protocol was approved Ministry of Agriculture of the Czech Republic (reference number: 55187/2016-MZE-17214)

## Consent for publication

Not applicable.

## List of abbreviations

GSCs: Germ stem cells
PGCs: Primordial germ cells
PBS: Phosphate buffered saline
FBS: Foetal bovine serum
DPF: Days post fertilisation
WPF: Weeks post fertilisation
SEM: Scanning electron microscopy
TEM: Transmission electron microscopy
GC: Germline chimera
CC: Common carp
GF: Goldfish

